# Direct and transgenerational effects of an experimental heat wave on early life stages in a freshwater snail

**DOI:** 10.1101/449777

**Authors:** Katja Leicht, Otto Seppälä

**Affiliations:** Eawag, Swiss Federal Institute of Aquatic Science and Technology, 8600 Dübendorf, Switzerland; Department of Biological and Environmental Science, PO Box 35, 40014 University of Jyväskylä, Finland; ETH Zürich, Institute of Integrative Biology (IBZ), 8092 Zürich, Switzerland

**Keywords:** climate change, environmental stress, global warming, *Lymnaea stagnalis*, maternal effects

## Abstract

Global climate change imposes a serious threat to natural populations of many species. Estimates of the effects of climate change-mediated environmental stresses are, however, often based only on their direct effects on organisms, and neglect the potential transgenerational effects. We investigated whether high temperature (i.e. an experimental heat wave) that is known to reduce the performance of adult *Lymnaea stagnalis* snails affects their offspring through maternal effects. Specifically, we tested whether eggs and hatched juveniles are affected by maternal thermal environment, and how strong these effects are compared with direct effects of temperature on offspring. We examined the effect of maternal thermal environment (15°C versus 25°C) on per offspring investment (egg size), and the role of both maternal and offspring thermal environments (15°C versus 25°C) on hatching success and developmental time of eggs, offspring survival after hatching, and hatchling size at the age of five weeks. Exposure of mothers to high temperature increased hatching success of eggs, and also made the onset of hatching earlier. However, high maternal temperature reduced the survival and the final size of hatched juveniles. Direct effects of high temperature on offspring survival were negative (both eggs and hatchlings), but increased the developmental rate and growth of those eggs and hatchlings that survived. Interestingly, the magnitude of transgenerational effects of high temperature on hatching success of eggs and hatchling survival were similar to its direct effects. This indicates that heat waves can affect natural populations through transgenerational effects and that the magnitude of such effects can be equally strong to the direct effects of temperature, although this depends on the trait considered. Our results highlight the importance of considering the transgenerational effects of climate warming when estimating its effects in the wild.

## Introduction

Owing to global climate change, the average temperatures at the Earth’s surface, as well as the frequency and severity of extreme weather events such as summer heat waves, are increasing (Easterling et al. 2000; Karl & Trenberth 2003; Meehl & Tebaldi 2004; Kirtman et al. 2013). These changes can have strong effects on organisms that escalate to higher levels of biological organization such as populations and communities (Walther et al. 2002; Parmesan & Yohe 2003; Walther 2010). Especially extreme weather events can dramatically influence population dynamics, species abundance, and species interactions (e.g. Easterling et al. 2000; Bruno et al. 2007; Hance et al. 2007). However, environmental conditions can not only influence the fitness of the individuals exposed to them but also the fitness of their offspring through transgenerational maternal and/or paternal effects (reviewed in Bernardo 1996; Mousseau & Fox 1998b). Hence, for understanding the effects of climate change on natural populations, studies examining such transgenerational effects are needed.

In particular, transgenerational maternal effects after exposure to environmental stress can significantly alter offspring performance (e.g. Silbermann & Tatar 2000; Mitchell & Read 2005; Janhunen, Piironen & Peuhkuri 2010). Such effects can result from the reduced physiological condition of the mother that limits the total amount of resources it invests in reproduction (Tessier et al. 1983; Steer et al. 2004), and/or exposure of offspring to hormones produced by the mother (McCormick 1999; Groothuis & Schwabl 2008). Maternal effects could also take place via altered resource allocation between reproduction and other traits depending on the environmental conditions the mother experiences. Challenging environmental conditions may, for example, reduce resource allocation to produced offspring to sustain self-maintenance or increase per offspring investment when the reproductive value of individuals is changed so that investment in current reproduction increases at the expense of future reproduction (Fisher 1930; Williams 1966). Furthermore, maternal effects can be adaptations to prepare offspring for the future conditions they are about to encounter (e.g. herbivory, parasitism, pollution; Agrawal 2002; Moret 2006; Marshall 2008).

To understand the consequences of such transgenerational effects in the context of climate change, it is essential to estimate their direction and magnitude compared with the direct effects of the same environmental factors. Maternal effects are typically strongest in the early stages of organisms’ life histories (Mousseau & Dingle 1991; Heath, Fox & Heath 1999; Pettay et al. 2008), but such stages are often also highly susceptible to the direct effects of environmental variation (e.g. Jang 1991; Zhang et al. 2015; Klockmann, Günter & Fischer 2017). For example, temperature determines the development of eggs and juveniles by altering their metabolic and physiological processes in many species (Gillooly et al. 2001; Person-Le Ruyet et al. 2004; Zuo et al. 2012). The high temperature, in particular, can impose a serious challenge by reducing the hatching success of eggs and early survival of hatched offspring (Janhunen et al. 2010; Zhang et al. 2015; Klockmann et al. 2017). Despite high interest on transgenerational effects of climate change in natural populations (reviewed in Donelson et al. 2018), their relative importance compared with direct effects of the same environmental factors is, however, often overlooked (but see Burgess & Marshall 2011; Parker et al. 2012; Salinas & Munch 2012; Shama et al. 2014; Wadgymar, Mactavish & Anderson 2018).

Here, we tested whether high temperature as it can occur during heat waves has transgenerational effects on offspring performance, which traits they affect, and how strong they are compared with direct effects of high temperature in the freshwater snail *Lymnaea stagnalis* L. (Gastropoda: Pulmonata). In this species, exposure of adult individuals to high temperature (≥ 25°C) initially increases growth and reproduction, but prolonged exposure (one week or longer) ceases reproductive rate and reduces immune function (Seppälä & Jokela 2011; Leicht, Jokela & Seppälä 2013). This indicates that high temperature is physiologically challenging and has strong negative effects on adult snails. We estimated the effect of maternal thermal environment (15°C versus 25°C) on per offspring investment by adult snails (egg size), and the role of both maternal and offspring thermal environments on offspring performance (hatching success and developmental time of eggs, survival of hatched offspring, offspring size at the age of five weeks) using a full-factorial design. We found that high temperature affected offspring performance both directly and through maternal effects. The relative importance and the direction of these effects varied among traits, and the magnitude of maternal effects was equally strong to direct effects in some of the examined traits. This highlights the importance of considering transgenerational effects when estimating the consequences of climate change in natural populations.

## Methods

### Experimental animals

The snails used in this study came from a laboratory stock population (F_4_ generation) originating from a pond in Zurich, Switzerland (47°22’05’’N, 8°34’41’’E). The summer water temperature in ponds typically remains low (< 16°C) in this region, although it depends on pond hydrology (T. Salo, unpublished data). However, during heat waves, water temperature can rapidly increase to 20–30°C and remain high for over two weeks (T. Salo, unpublished data). We started the stock population using 45 adult snails collected from the pond. Since *L. stagnalis* prefers outcrossing (Puurtinen et al. 2007; Nakadera et al. 2017), often engages in multiple matings (Nakadera et al. 2017), and can store sperm from those matings for over two months (Nakadera, Blom & Koene 2014), the stock population can be expected to reflect the genetic variation in the source population well. We maintained the stock population in the approximate size of 400 individuals at 15 ± 2°C (control temperature used in the experiment; see the section about experimental design below) for two years before the study (see Leicht, Seppälä & Seppälä 2017).

We haphazardly collected 113 adult snails from the stock population and used them as a maternal generation in the experiment. We placed the snails individually in 2 dl perforated plastic cups sunk into a water bath (aged tap water at 15 ± 1°C) that was connected to a biological filter. We used a water bath to provide maximal water quality for snails. This minimises the growth of microorganisms in water that activate snail immune function (Seppälä & Leicht 2013). This is important because immune challenge could potentially alter snail reproductive strategy and/or affect the quality of produced offspring. We fed the snails with fresh lettuce ad libitum and maintained them under these conditions for three days prior to the experiment to acclimate them to the maintenance conditions. Because *L. stagnalis* snails can reproduce through self-fertilization as well as through outcrossing using allosperm they have stored from previous matings (Cain 1956; Nakadera et al. 2014), experimental snails did not need a mating partner to oviposit eggs under the used conditions.

### Experimental design

#### Maternal treatments

At the beginning of the experiment, we randomly assigned the snails used as a maternal generation (see the previous section) into two temperature treatments [15 ± 1°C (54 snails), 25 ± 1°C (59 snails); Fig. 1]. We used 25°C as a high (i.e. heat wave) temperature as it reduces immune defence and life history traits in adult snails (Seppälä et al. 2011; Leicht et al. 2013), lies above the thermal optimum for development and growth of juvenile snails (Vaughn 1953), and occurs intermittently in habitats of snails during hot summers (T. Salo, unpublished data). We chose 15°C as a control temperature as it is close to the thermal optimum of *L. stagnalis* (Vaughn 1953) and common in ponds (T. Salo, unpublished data). Note that we assigned more individuals into the high-temperature treatment because we expected increased mortality in those snails. We transferred the snails to their treatment temperatures in cups filled with aged tap water at 15°C. This allowed a slow change (over 10 h) to the target temperature for snails assigned to the high-temperature treatment. We then transferred the snails into perforated plastic cups (2 dl) sunk into similar water baths as above, and exposed them to their respective temperature treatments for seven days. At 15°C, 51 of these snails survived, and of those 25 oviposited eggs. At 25°C, 39 snails survived, and 34 of them reproduced. We did not measure the number of oviposited eggs in this experiment as the effect of temperature on snail fecundity has been described in detail in earlier studies (Leicht et al. 2013; 2017).

**Figure 1.**
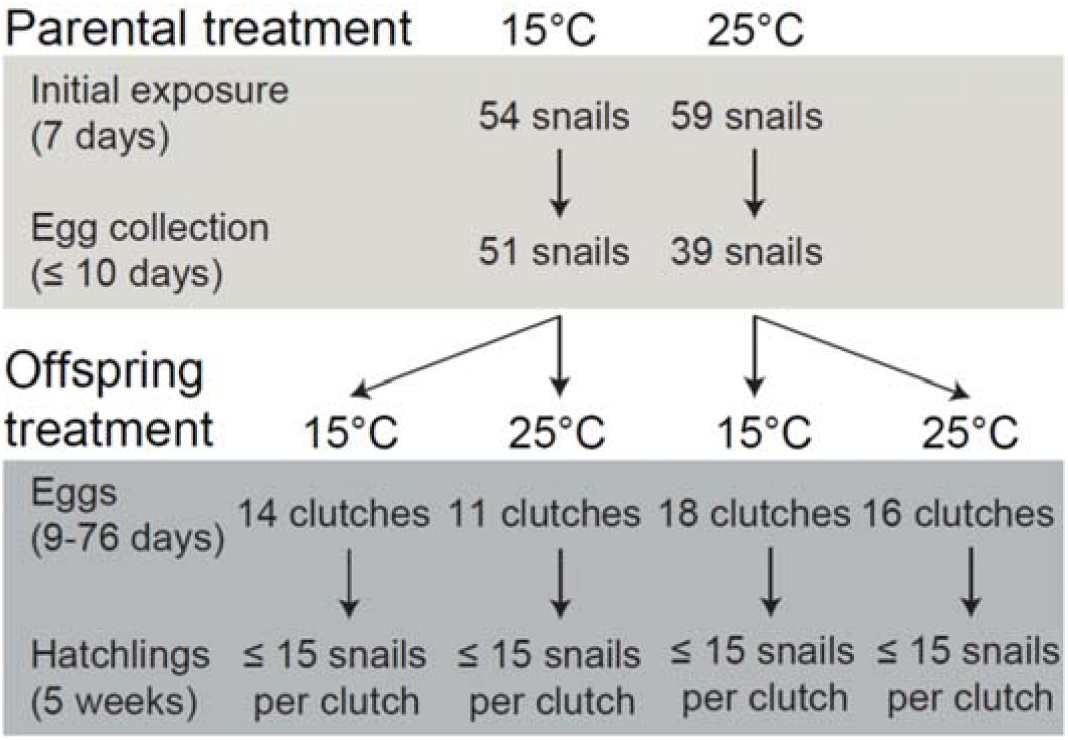
Summary of the full-factorial experimental design used to examine direct and transgenerational effects of exposure of *L. stagnalis* snails to two temperatures (benign: 15°C; heat wave: 25°C). One egg clutch per adult snail was used in offspring treatments. When available, 15 hatchlings per egg clutch were used. When fewer individuals were available, they were all used.

After the initial seven-day exposure to temperature treatments, we removed all egg clutches oviposited by the snails from the cups and continued maintaining the snails under the same experimental conditions. During the following ten days, we checked the cups twice a day for new clutches to be collected for the next step of the experiment (see the next section). This procedure ensured first, that the snails were exposed to their respective temperature treatments long enough to induce strong effects on their performance before the clutches were collected (see Leicht et al. 2013), and second, that the collected clutches were exposed to the maternal temperature treatments only briefly. From each snail that oviposited (same individuals as above reproduced), we collected the first clutch containing more than 20 eggs, or if only smaller clutches were produced, the largest clutch. We placed each collected clutch on a millimetre paper and photographed it from 10 cm above with a Fujifilm FinePix F30 digital camera (scene mode: close up, focal length: 35 mm, aperture: F/2.8, shutter speed: 1/85, sensitivity: ISO-200, image size: 2848 × 2136 pixels, focus mode: auto focus). From the digital images, we counted the eggs in each clutch. Furthermore, we measured the two-dimensional area (mm^2^) of five randomly chosen eggs in each clutch from the digital images using ImageJ software (ImageJ 1.42q, Wayne Rasband, National Institute of Health, USA). After photographing, we placed the clutches individually into plastic cups with 0.4 dl of aged tap water to be transferred to the next step of the experiment (see the next section). It is important to note that the time different snails needed for ovipositing after the initial exposure period varied between one and ten days, which may have affected the thermal challenge imposed to them as well as their offspring. However, to our knowledge, oviposition cannot be induced artificially in this species.

#### Offspring treatments

We used a full-factorial design to expose egg clutches produced in both maternal temperature treatments (see the previous section) to two offspring temperature treatments (15°C, 25°C; Fig. 1). In each maternal temperature treatment, we randomly assigned some of the oviposited egg clutches to remain at the same temperature where they were produced and transferred the rest of the clutches to the other temperature. The number of egg clutches per treatment combination varied between 11-18, which was because of unequal mortality and probability to reproduce in parental snails in different treatments (see the previous section). We slowly warmed up or cooled down the clutches that were transferred to a different temperature as described above to avoid a sudden change between temperatures. After that, we checked the clutches daily, counted the number of hatched snails, and removed the hatchlings from the cups. When possible, we placed 15 hatchlings from each clutch individually in plastic cups filled with 0.4 dl of aged tap water. We used all hatchlings when less than 15 individuals were available. We fed the snails with Spirulina ad libitum and changed the water in the cups twice a week. We reared the isolated hatchlings for five weeks and measured their survival and shell length to the nearest 0.1 mm using a digital calliper at the end of the experiment.

### Statistical analyses

We analysed the effect of temperature on the survival of adult snails during the initial exposure to experimental treatments (a seven-day period before the collection of egg clutches started) using a generalized linear model. We used the status of snails (survived, died) as a binomial response variable (logit link function) and temperature treatment as a fixed factor. Additionally, we analysed variation in snail reproductive status (oviposited, did not oviposit) using a similar generalised linear model. We analysed the effect of temperature on the size of produced eggs (ln transformed to homogenize error variance) using a mixed-model analysis of variance (mixed-model ANOVA). In the analysis, we used a model with maternal temperature treatment as a fixed and the clutch each egg originated from as a random factor (nested within maternal temperature).

To estimate the effects of maternal and offspring temperature on offspring performance, we first analysed the variation in hatching success of eggs using a generalized linear model. In the analysis, we used the proportion of eggs that hatched from each clutch as a binomial response variable (logit link function), and maternal temperature treatment and offspring temperature treatment as fixed factors. We included the interaction term between the factors into the model. Less than three snails hatched from two clutches that were both produced and maintained at 25°C. We excluded these individuals from all the further analyses as they would not provide sufficient replication within those clutches.

After that, we calculated the developmental time until hatching for each egg as the difference between the date the clutch was oviposited and the hatching date. We then analysed the effects of temperature on developmental time using a multivariate analysis of variance (MANOVA, with Pillai’s trace test statistic for unequal sample sizes). We used the onset of hatching (i.e. the first hatching day; square-root transformed to homogenize error variance), median developmental time (we used the median rather than the mean as the distribution of hatching time within the clutches was skewed), and the end of hatching (i.e. the last hatching day; ln transformed to homogenize error variance) for each clutch as response variables. We used maternal temperature treatment and offspring temperature treatment as fixed factors in the analysis and included the interaction term between them into the model. Since the MANOVA indicated effects of temperature on developmental time (see the results section), we conducted separate ANOVAs using a similar model as above for the different parameters of developmental time to investigate whether their responses to temperature were different.

We analysed the variation in the survival of hatched offspring during the experiment using a generalized linear mixed-effects model with the status of snails (survived, died) as a binomial response variable (logit link function). We used maternal temperature treatment and offspring temperature treatment as fixed factors, and egg clutch each offspring originated from as a random factor (nested within the interaction between maternal temperature and offspring temperature). We included the interaction term between maternal and offspring temperature into the model. From the offspring that survived until the end of the experiment, we analysed the variation in size using a mixed-model ANOVA with shell length (square-root transformed to homogenize error variance) as a response variable, and a similar model as for survival. Survival and/or size could not be measured from 30 juvenile snails (3.5% of all individuals) because of human errors. We excluded these snails from the data. The assumptions of all the above analyses were fulfilled, and we performed them using IBM SPSS Statistics Version 23.0 software (Armonk, NY: IBM Corp.).

## Results

During the initial exposure of adult snails to different temperature treatments (i.e. a seven-day period before the collection of egg clutches started), the survival of snails exposed to 25°C was reduced (estimated marginal mean ± SE: 66.1 ± 6.2%) compared with snails exposed to 15°C (estimated marginal mean ± SE: 94.4 ± 3.1%; generalized linear model: Wald Chi-Square = 10.940, d.f. = 1, p = 0.001). In those snails that survived, the probability of reproducing was higher at 25°C (estimated marginal mean ± SE: 87.2 ± 5.4%) than at 15°C (estimated marginal mean ± SE: 49.0 ± 7.0%; generalized linear model: Wald Chi-Square = 12.429, d.f. = 1, p < 0.001). Note that the same individuals that reproduced during this initial exposure period also produced the eggs that were exposed to offspring temperature treatments.

Eggs oviposited by snails at 25°C were smaller than those oviposited at 15°C (two-dimensional area; estimated marginal mean ± SE: 25°C: 1.11 ± 0.01 mm^2^; 15°C: 1.30 ± 0.01 mm^2^; ANOVA: F_1,57_ = 26.275, p < 0.001). Hatching success of eggs was affected by both the maternal temperature treatment and the offspring temperature treatment, and these effects were independent of each other (Table 1, Fig. 2). High maternal temperature increased hatching success by 9.0% whereas high offspring temperature reduced it by 7.5% (Fig. 2). Maternal and offspring temperature also affected the developmental time of eggs, and these effects were independent of each other (MANOVA, maternal temperature treatment: Pillai’s trace = 0.310, F_3,51_ = 7.654, p < 0.001; offspring temperature treatment: Pillai’s trace = 0.918, F_3,51_ = 191.413, p < 0.001; maternal temperature treatment × offspring temperature treatment: Pillai’s trace = 0.046, F_3,51_ = 0.827, p = 0.485). The effects of temperature on developmental time were first because offspring started to hatch 12.3% earlier when mothers had been exposed to 25°C (ANOVA: F_1,53_ = 15.806, p < 0.001; Fig. 3a). Second, the onset, median, and the end of hatching were earlier when offspring were maintained at 25°C (first day of hatching: 55.1% reduction; ANOVA: F_1,53_ = 571.961, p < 0.001; median developmental time: 43.8% reduction; ANOVA: F_1,53_ = 189.817, p < 0.001; last day of hatching: 43.8% reduction; ANOVA: F_1,53_ = 62.002, p < 0.001; Fig. 3).

**Table 1.**
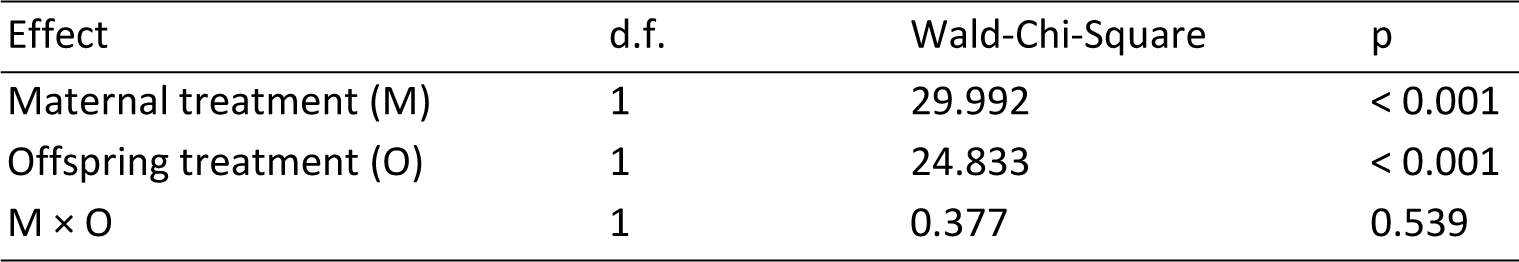
Generalized linear model for the hatching success of *L. stagnalis* eggs (proportion of eggs that hatched per clutch) by maternal temperature treatment (15°C, 25°C), offspring temperature treatment (15°C, 25°C), and their interaction.

**Figure 2.**
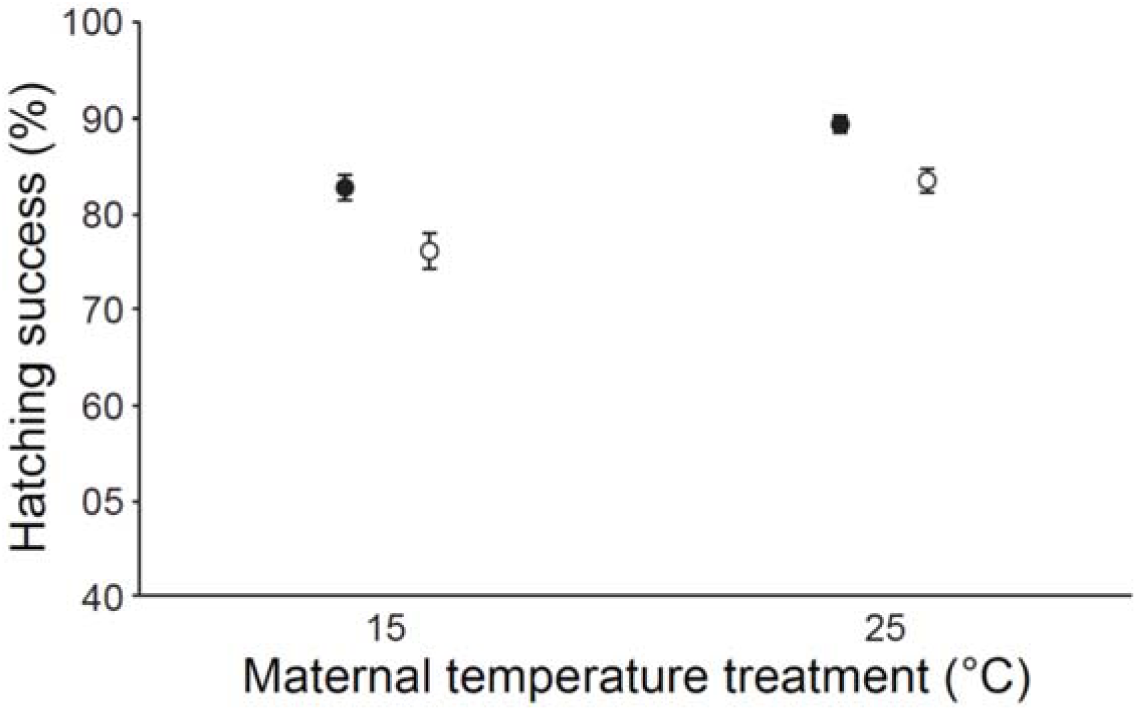
Hatching success of eggs [proportion of eggs that hatched (%; mean ± SE)] for egg clutches produced at different maternal temperature treatments (15°C, 25°C) and maintained at 15°C (black circles) or at 25°C (white circles).

**Figure 3.**
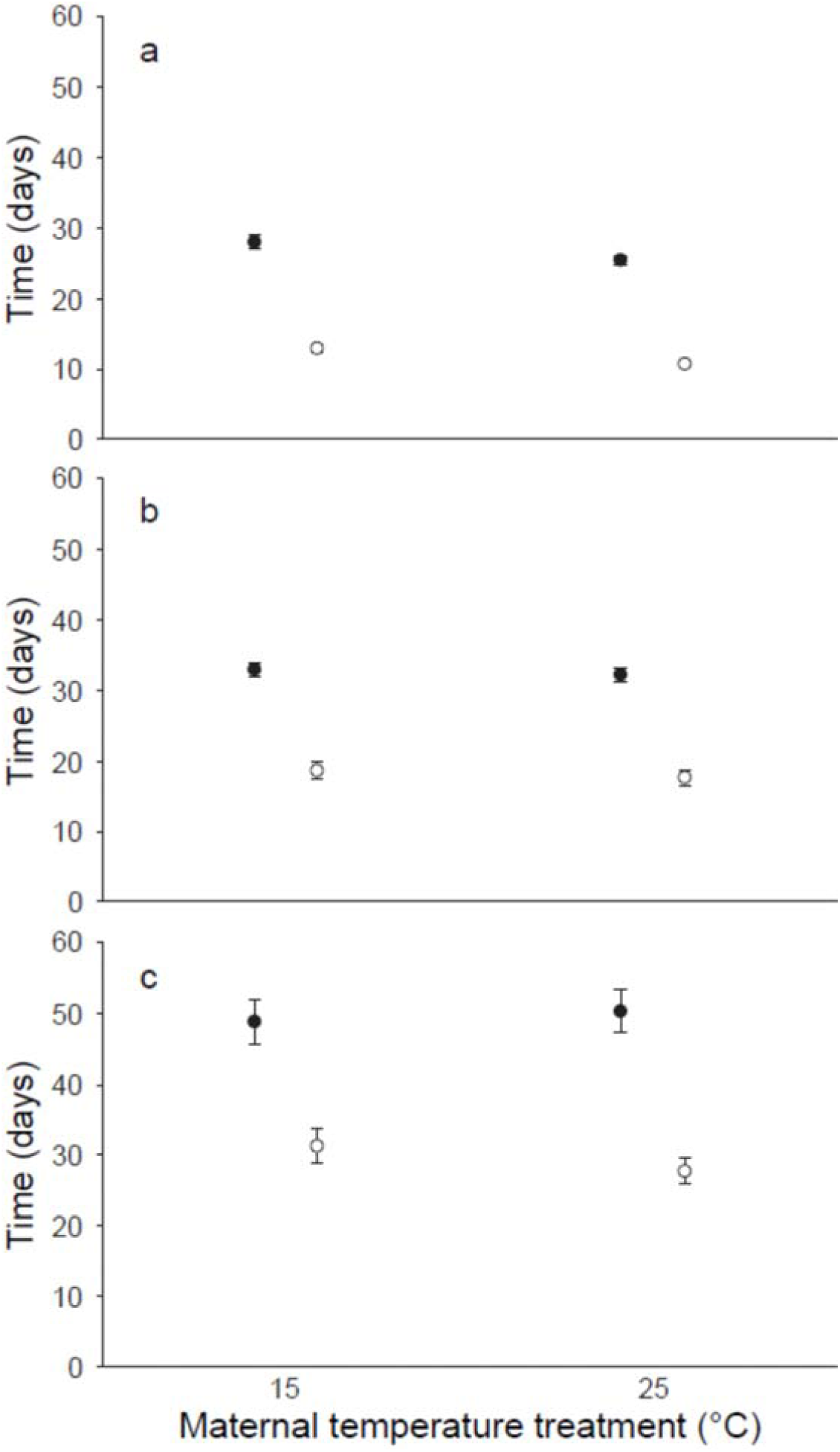
Developmental time of eggs presented using the mean ± SE of (a) onset of hatching, (b) median developmental time, and (c) end of hatching for egg clutches produced at different maternal temperature treatments (15°C, 25°C) and maintained at 15°C (black circles) or at 25°C (white circles).

High maternal temperature reduced offspring survival by 17.8%, and this effect was independent of the temperature offspring were exposed to (Table 2, Fig. 4a). When offspring were maintained at 25°C, they showed a tendency towards lower survival (Table 2, Fig. 4a). Temperature affected the size of offspring so that high maternal temperature and low offspring temperature reduced shell length at the end of the experiment (Table 3, Fig. 4b).

**Table 2.** Generalized linear mixed-effects model for the survival of juvenile *L. stagnalis* snails during the experiment (survived/died) by maternal temperature treatment (15°C, 25°C), offspring temperature treatment (15°C, 25°C), their interaction, and the egg clutch each individual originated from.

| Effect | d.f. | F | Estimate | SD | Z | p |
| --- | --- | --- | --- | --- | --- | --- |
| Maternal treatment (M) | 1 | 4.452 |  |  |  | 0.035 |
| Offspring treatment (O) | 1 | 3.588 |  |  |  | 0.059 |
| M × O | 1 | 1.587 |  |  |  | 0.208 |
| Egg clutch (M × O) |  |  | 0.582 | 0.184 | 3.165 | 0.002 |

**Figure 4.**
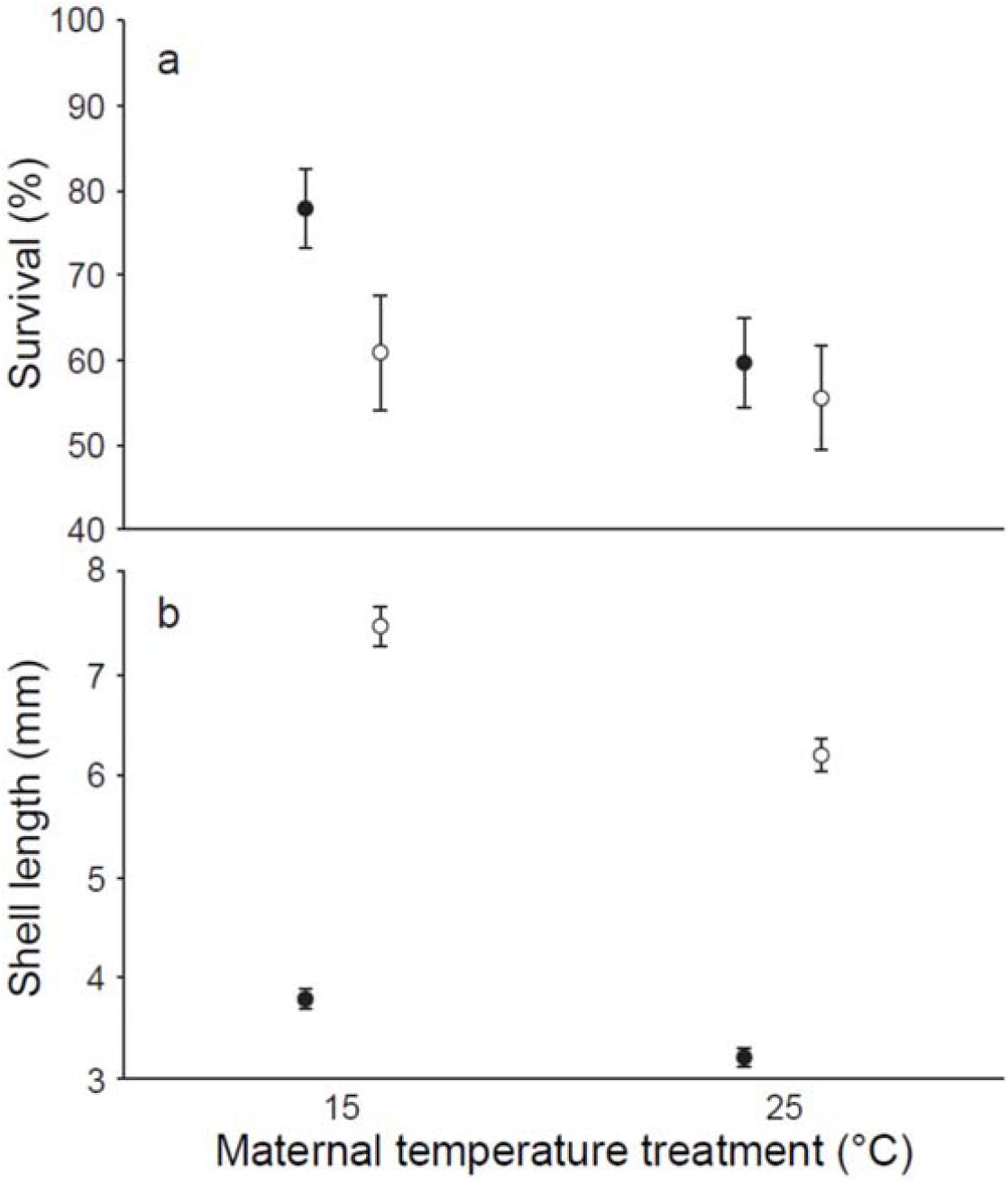
(a) Proportion (%; estimated marginal mean ± SE) of offspring that survived until the end of the experiment, and (b) shell length (mm; estimated marginal mean ± SE) of those snails at the end of the experiment when produced at different maternal temperature treatments (15°C, 25°C) and maintained at 15°C (black circles) or at 25°C (white circles).

**Table 3.**
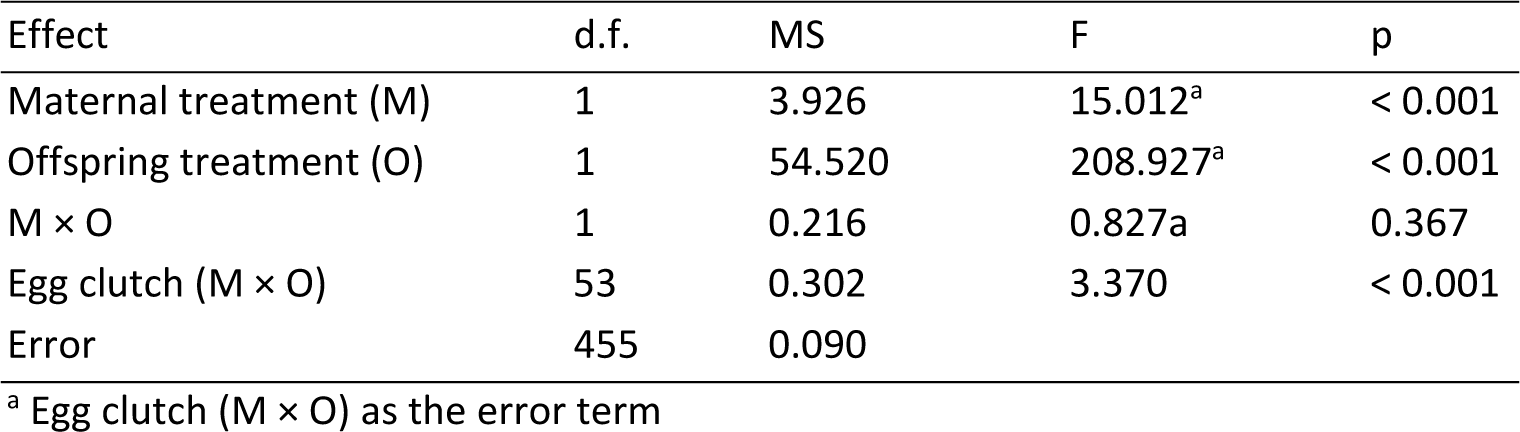
Mixed-model analysis of variance for the shell length of juvenile *L. stagnalis* snails at the end of the experiment by maternal temperature treatment (15°C, 25°C), offspring temperature treatment (15°C, 25°C), their interaction, and the clutch each individual originated from.

| Effect | d.f. | MS | F | p |
| --- | --- | --- | --- | --- |
| Maternal treatment (M) | 1 | 3.926 | 15.012<sup>a</sup> | < 0.001 |
| Offspring treatment (O) | 1 | 54.520 | 208.927<sup>a</sup> | < 0.001 |
| M × O | 1 | 0.216 | 0.827a | 0.367 |
| Egg clutch (M × O) | 53 | 0.302 | 3.370 | < 0.001 |
| Error | 455 | 0.090 |  |  |

<sup>a</sup> Egg clutch (M × O) as the error term

## Discussion

Exposure to an experimental heat wave affected eggs and hatchlings of *L. stagnalis* snails both directly and via transgenerational maternal effects. Direct effects of high temperature were both negative and positive by reducing the hatching success of eggs, but by increasing their development rate and the subsequent growth of those individuals that hatched. The magnitude and the direction of transgenerational effects when compared with direct effects varied among examined traits. In general, high maternal temperature benefitted offspring at very early life stages (hatching success and the onset of hatching), but reduced performance at later stages (survival and final size of hatched snails). Hence, the potential adaptive value of responding to high temperature by maternal effects may be limited only to very early life stages. Interestingly, the magnitude of transgenerational effects on hatching success and survival of offspring were similar to the direct effects of high temperature, although in the case of hatching success the direction of the effects differed. These findings indicate that heat waves can not only impact natural populations through transgenerational effects but that the magnitude of those effects can be equally strong to the direct within generation effects of high temperature. The relative strength of the direct and transgenerational effects, however, depends on the considered trait.

It is important to note that because all parental snails that we initially exposed to different temperatures did not survive or reproduce, potential differences between the temperature treatments in offspring generation could arise from selection in the parental population. This could be, for instance, if weak snails could not survive under environmental stress. Similarly, not all offspring hatched or they died before the end of the experiment. While we cannot exclude either estimate the potential role of selection in our experiment, it seems unlikely that selection could explain our results on transgenerational effects. This is because the longest lasting effects of high maternal temperature on offspring (i.e. hatchling survival and size at the end of the experiment) were negative. If only high-quality individuals were able to survive and reproduce at the high temperature this should lead to the opposite result. Because high temperature had a direct negative effect on the hatching of juvenile snails, their increased size at the end of the experiment could be not only due to phenotypic plasticity but also because of selection.

### Effects of temperature on eggs

Mothers exposed to high temperature produced smaller eggs that had higher hatching success compared with snails that oviposited at the benign temperature (14.6% reduction in size, 9.0% increase in hatching success). Thus, exposure of the maternal generation to high temperature can increase the quality of produced eggs despite their smaller size. The high maternal temperature made the onset of hatching earlier by 2.5 days (12.3% reduction). The magnitude of the direct effect of high temperature on the hatching success of eggs was similar to the observed transgenerational effect, but negative (7.5% reduction). The direct effect of high temperature on the development of eggs was strong by shortening the median developmental time by approximately two weeks (43.8% reductions). Faster development and earlier hatching could reduce the risk of eggs being exposed to natural enemies such as predators (see Warkentin 1995; Chivers et al. 2001), and it is suggested to increase future survival of offspring (Arcese & Smith 1985; Warner & Shine 2007) as well as allow earlier maturation and increased reproductive output (Uller & Olsson 2010). However, early hatching may also bring disadvantages by leading to less well-developed offspring (e.g. Warkentin 1999; Buckley, Michael & Irschick 2005).

Large egg size is often beneficial for offspring by increasing their fitness (Hutchings 1991; Fox 1994; Krist 2011). This is likely to be because large eggs can provide more energy and nutrients for developing embryos (reviewed in Williams 1994). In aquatic organisms, however, reduced egg size may be beneficial by ensuring oxygen supply to embryos when the oxygen concentration in water decreases under high temperature (Woods 1999; Moran & Woods 2007). Therefore, egg size might not be a good indicator of egg quality. The observed transgenerational effect of high temperature on the development of eggs (developmental time and hatching success) may be due to two non-exclusive mechanisms. First, high temperature can increase the metabolic rate of adults that allow the production of high-quality eggs (see Jann & Ward 1999; Saino et al. 2004). Second, it may increase resource allocation towards reproduction rather than other traits (e.g. growth) when the residual reproductive value of individuals decreases, for example, due to increased mortality (Fisher 1930; Williams 1966). The effect of maternal temperature on the developmental time of eggs was, however, limited to the onset of hatching. This indicates that the possibly increased investment on oviposited eggs may be rapidly depleted. On the other hand, smaller eggs are expected to develop more slowly (Levitan 2000). Therefore, it is possible that high maternal investment and small egg size overrode each other’s effects so that no net change in developmental time of eggs could be detected. The direct effect of temperature on hatching success of eggs is likely to be due to the high sensitivity of mollusc embryos to high temperature (Vaughn 1953), which leads to mortality in several taxa probably due to denaturation of proteins (reviewed in Pepin 1991; Noble, Stenhouse & Schwanz 2018). The direct effect of temperature on the developmental time of eggs was possibly because temperature determines the speed of biochemical processes of the developing embryos (e.g. García-Guerrero, Villarreal & Racotta 2003; Sibert, Ouellet & Brêthes 2004).

### Effects of temperature on hatchlings

Exposure of mothers to high temperature reduced the probability of offspring to survive until the age of five weeks (17.8% reduction compared with 15°C). High maternal temperature also reduced the size offspring reached (28.9% reduction). High temperature showed a tendency to have a negative direct effect on offspring survival (16.1% reduction). However, offspring that survived grew larger at high temperature (97.4% increase in size). Hence, high maternal temperature reduced offspring performance in the examined traits while the direct effects of high temperature were both positive and negative. Reduced size of offspring due to high maternal temperature may lead to delayed maturity as well as reduced mating success and fecundity (reviewed in Clutton-Brock 1988), and also increase susceptibility to predators (e.g. Janzen 1993; Craig et al. 2006). Instead, the direct effect of high temperature can benefit those individuals that are able to survive under such conditions.

Reduced offspring size when mothers experience high temperature is found across a wide range of animal taxa (reviewed in Atkinson et al. 2001). The reason for this is not yet clear and may either be an adaptation to maximize mother’s life time fitness (Yampolsky & Scheiner 1996) or due to physiological constraints under such conditions (Blanckenhorn 2000). The direct effect of high temperature on offspring survival may be due to temperature-induced changes in, for instance, protein structures and/or membrane fluidity (reviewed in Pörtner, Lucassen & Storch 2005), which can lead to body malfunctions and increased mortality. On the other hand, high temperature fastens metabolic rate and can increase the growth of organisms (e.g. Iguchi & Ikeda 2005; Salo et al. 2017). This, however, could also lead to faster use of energetic reserves that may reduce survival.

### General conclusions

Our finding that the potential adaptive value of responding to high temperature by maternal effects was limited to very early life stages (i.e. eggs) is in line with earlier research (Mousseau et al. 1991; Heath et al. 1999; Pettay et al. 2008). Instead, the result of equally strong direct and transgenerational effects of exposure to high temperature on some of the examined traits contradict earlier studies that have examined their relative importance in determining offspring performance and physiology (e.g. Groeters & Dingle 1988; Huey et al. 1995; Steigenga & Fischer 2007; Burgess et al. 2011; Salinas et al. 2012; Shama et al. 2014). In those studies, transgenerational effects of temperature are typically reported to be weak compared with its direct effects. To our knowledge, transgenerational effects of environmental change have been found to be strong compared with its direct effects in a climate change context only in germination probability of a perennial forb, *Boechera stricta*, when wintering conditions are manipulated (Wadgymar et al. 2018). Together with that finding, our results indicate that climate change-mediated environmental changes can affect natural populations through transgenerational effects and that these effects may be as strong as or even stronger than the direct effects of environmental change.

Despite high interest in transgenerational effects of climate change-mediated environmental change on organisms (Donelson et al. 2018), many studies have not tested their relative importance compared with direct within generation effects. This is because earlier studies have focused, for example, on testing whether negative effects of environmental change are reduced if parents have experienced the same environmental conditions (e.g. Donelson et al. 2012; Miller et al. 2012). Testing this does not necessarily require a full-factorial design that is needed for examining the relative importance of direct and transgenerational effects. The alternative approach is relevant in systems where the environment changes gradually and relatively slowly to a predicted direction (e.g. in oceans). Such studies would, however, not be realistic in terrestrial and freshwater systems that experience high and rapid fluctuations in several environmental conditions owing to extreme weather events such as heat waves. In marine species, exposing parents to altered environmental conditions has been found to reduce the negative effects of increased temperature and CO2-level on offspring (Donelson et al. 2012; Miller et al. 2012; Shama et al. 2014). In our study, none of the observed direct effects of temperature depended on the maternal environment, which was indicated by the lack of interactive effects between temperature treatments. This may be due to higher unpredictability of extreme weather events in freshwater systems compared with marine environments that could limit the ability of such adaptive maternal effects to evolve (see Mousseau & Fox 1998a).

## Data accessibility

Data are available online: DOI: 10.5281/zenodo.2600811

## Acknowledgements

We thank K. Räsänen and J. Jokela for constructive comments on the manuscript. The study was funded by the Biological Interactions Doctoral Program (BIOINT) to KL and the Emil Aaltonen Foundation and the Swiss National Science Foundation (grant 31003A 140876 and 31003A 169531) to OS. This preprint has been reviewed and recommended by Peer Community In Ecology (https://dx.doi.org/10.24072/pci.ecology.100015).

